# Functional characterization of the strawberry calcium-dependent protein kinase FvCPK28

**DOI:** 10.64898/2026.10.26.762000

**Authors:** Márcia Gonçalves-Dias, Jacqueline Monaghan

**Affiliations:** Department of Biology, Queen’s University, Kingston, ON, Canada; Biological Sciences Department, Luiz de Queiroz College of Agriculture, University of São Paulo, Piracicaba, Brazil

**Keywords:** CPK28, BIK1, pattern-triggered immunity, immune homeostasis, Rosaceae, *Fragaria vesca*

## Abstract

Calcium-dependent protein kinases (CDPKs) translate cellular calcium signals into phosphorylation events that regulate plant development and immunity. In *Arabidopsis thaliana*, CPK28 restrains pattern-triggered immunity by limiting the accumulation of the receptor-like cytoplasmic kinase BOTRYTIS-INDUCED KINASE 1 (BIK1), and contributes to roles in development such as stem elongation. CPK28 orthologs have been described in several crops, but not in Rosaceae, a family that includes economically important fruits such as strawberry, apple, peach, and raspberry. Phylogenetic analysis of CDPKs and CDPK-related kinases (CRKs) in four Rosaceae species and Arabidopsis identified CPK28 orthologs forming a clade with Arabidopsis CPK28. Using the woodland strawberry (*Fragaria vesca*) ortholog, FvCPK28, as a case study, we show that FvCPK28 is a calcium-dependent kinase that phosphorylates both Arabidopsis AtBIK1 and a strawberry FvBIK1 homolog *in vitro* and associates with both proteins *in planta*. When expressed in the Arabidopsis *cpk28-1* mutant, FvCPK28 restored stem elongation and immune responses to wild-type levels. These results show that the strawberry ortholog retains the biochemical and biological properties of CPK28. This work extends the functional study of CPK28 to the Rosaceae and identifies FvCPK28 as a candidate for modulating the balance between growth and defense in strawberry.

**Main Conclusion:** Strawberry FvCPK28 retains the biochemical and genetic properties of Arabidopsis CPK28, functioning as a Ca^2+^-dependent kinase that associates with and phosphorylates AtBIK1 and complements the *cpk28-1* mutant.

## Introduction

Plants defend themselves against pathogens using a multilayered immune system. At the cell surface, pattern recognition receptors (PRRs) perceive conserved microbe- and damage-associated molecular patterns (MAMPs/DAMPs), activating pattern-triggered immunity (PTI). Upon ligand perception, PRRs such as FLAGELLIN SENSING 2 (FLS2) associate with the co-receptor BRASSINOSTEROID INSENSITIVE 1-ASSOCIATED RECEPTOR KINASE 1 (BAK1) (Heese et al. 2007; Chinchilla et al. 2007). Following PRR activation, there is an influx of calcium (Ca^2+^), an apoplastic reactive oxygen species (ROS) burst, and activation of mitogen-activated protein kinase (MAPK) cascades, which initiate signaling cascades that ultimately restrict pathogen growth (DeFalco and Zipfel 2021). Because uncontrolled or constitutive immune activation impairs growth and reduces fitness, the amplitude and duration of immune signaling must be tightly controlled (Huot et al. 2014).

A central node in this regulation is the receptor-like cytoplasmic kinase (RLCK) BOTRYTIS-INDUCED KINASE 1 (BIK1), which acts immediately downstream of multiple PRR complexes as a convergent, rate-limiting component of PTI (Lu et al. 2010; Zhang et al. 2010; DeFalco and Zipfel 2021). BIK1 belongs to RLCK subfamily VII, together with the related PBS1-LIKE (PBL) kinases. Its closest homolog, PBL1, also contributes to PTI (Zhang et al. 2010). Upon activation, BIK1 phosphorylates downstream targets, including the NADPH oxidase RESPIRATORY BURST OXIDASE HOMOLOG D (RBOHD) to drive the ROS burst (Li et al. 2014; Kadota et al. 2014), and Ca^2+^-permeable channels such as REDUCED HYPEROSMOLALITY-INDUCED Ca^2+^ INCREASE (OSCA1.3) (Thor et al. 2020). The activity and stability of BIK1 are controlled by multiple regulatory mechanisms (Monaghan et al. 2014; Wang et al. 2018a; Gonçalves Dias et al. 2022; Dou et al. 2025). The Ca^2+^-dependent protein kinase CPK28 is a key negative regulator of this module (Monaghan et al. 2014). CPK28 attenuates PTI by phosphorylating and promoting the activation of the E3 ubiquitin ligases PLANT U-BOX 25 (PUB25) and related E3 ligases PUB26, PUB22, and PUB24, which ubiquitinate BIK1 to promote its proteasomal turnover (Wang et al. 2018a; Dou et al. 2025). Loss of CPK28 function leads to BIK1 over-accumulation, heightened immune responses, and enhanced antibacterial resistance, establishing CPK28 as a brake that contributes to the amplitude of immune signaling (Monaghan et al. 2014; Wang et al. 2018a). Beyond immunity, CPK28 acts in development, where *Arabidopsis thaliana* (hereafter: Arabidopsis) *cpk28* mutants show an altered vegetative-to-reproductive transition and reduced stem elongation (Matschi et al. 2013). CPK28 also regulates ethylene biosynthesis and lignin deposition (Jin et al. 2017) and it is activated under cold stress to positively regulate the transcription factor NIN-LIKE PROTEIN 7 (NLP7) (Ding et al. 2022). In tomato, SlCPK28 contributes to thermotolerance by phosphorylating ascorbate peroxidase (Hu et al. 2021).

CPK28 belongs to the Ca^2+^-dependent protein kinase (CDPK/CPK) family, which is found in plants and some protists, including apicomplexans and oomycetes (Valmonte et al. 2014), and combines a serine/threonine kinase domain and Ca^2+^-binding EF-hands within a single polypeptide, linked by an autoinhibitory junction (Yip Delormel and Boudsocq 2019). This architecture allows CDPKs to translate cytosolic Ca^2+^ changes directly into phosphorylation events. The family is phylogenetically classified into five subgroups (I, II, III, IV, and CDPK-related kinases (CRKs)) (Yip Delormel and Boudsocq 2019). Notably, CRKs have been found only in plants, and their EF-hands are degenerate (Hrabak et al. 2003). Although CRKs are often not classified as *bona fide* CDPKs, we include them here as a fifth subgroup, given their close phylogenetic relationship to Subgroup IV.

Functional CPK28 orthologs that regulate immune signaling have been identified across diverse angiosperms, including rice (OsCPK4 and OsCPK18), cotton (GhCDPK28), and tomato (SlCPK28) (Xie et al. 2014; Wang et al. 2018b; Wu et al. 2021; Ding et al. 2022). Beyond angiosperms, a CPK28 ortholog from the liverwort *Marchantia polymorpha* (MpCPK28) complements the Arabidopsis *cpk28-1* mutant, although its role in immune homeostasis seems to have diverged in angiosperms as it is not conserved in *M. polymorpha* (Dou et al. 2026). By contrast, the economically important family Rosaceae, which includes strawberry, apple, pear, and raspberry, alongside numerous ornamentals, remains largely unexplored. These crops suffer substantial losses to pathogens (Da Costa Arruda et al. 2024), and enhancing their disease resistance remains a major breeding challenge.

Although the CDPK family has been catalogued in several Rosaceae genomes (Wei et al. 2016; Xiong et al. 2022), very few CDPKs have been functionally characterized in this family (Dong H, Wu C, Luo C, Wei M, Qu S, Wang S 2020; Crizel et al. 2020; Shi et al. 2025). Testing gene function and signaling pathways directly in Rosaceae species remains methodologically challenging. Cultivated strawberry (*Fragaria × ananassa*) is octoploid (Crizel et al. 2020), and apple (*Malus × domestica*) has long generation times and a highly duplicated genome, which frequently leads to genetic redundancy and functional divergence among paralogs (Lallemand et al. 2023). Although the diploid woodland strawberry (*Fragaria vesca*) has been used as the genetic model for the family due to its small genome and short life cycle (Shulaev et al. 2011), generating stable transgenic or genome-edited lines still requires several months (Zhou et al. 2018; Wilson et al. 2019), and *CPK28* loss-of-function mutants are not yet available. By contrast, the *Arabidopsis thaliana cpk28-1* mutant exhibits well-defined, highly reproducible developmental and immune phenotypes (Matschi et al. 2013; Monaghan et al. 2014), providing a robust, quantitative system to test whether a heterologous Rosaceae ortholog can replace AtCPK28 *in planta*.

Here, we identified CPK28 orthologs in Rosaceae through phylogenetic analysis of the CDPK/CRK family and characterized the *F. vesca* ortholog (FvH4_1g11780.1, hereafter FvCPK28). We show that FvCPK28 is an active Ca^2+^-dependent kinase that phosphorylates both Arabidopsis AtBIK1 and the strawberry homolog FvBIK1 *in vitro* and associates with both proteins *in planta*. Furthermore, transgenic expression of FvCPK28 complements the developmental and immune phenotypes characteristic of the Arabidopsis *cpk28-1* mutant. These findings suggest that FvCPK28 can functionally replace AtCPK28 and provide a starting point for investigating FvCPK28 function in Rosaceae.

## Materials and Methods

### Plant material and growth conditions

*Arabidopsis thaliana* Columbia-0 (Col-0) was used as the wild type. The T-DNA insertion mutant *cpk28-1* (GABI_523B08) and the complementation line *cpk28-1/35S:AtCPK28-YFP* were described previously (Matschi et al. 2013; Monaghan et al. 2014). The *cpk28-1/35S:FvCPK28-GFP* lines (#13 and #14) were generated in this study. Arabidopsis seeds were stratified at 4 °C for 2–3 days, then sown on soil or sterile Murashige and Skoog (MS) medium depending on the assay. For the oxidative burst assay, the genetic complementation lines were grown on soil (Sungro Sunshine Mix 1) for 4 weeks post-germination (wpg) in controlled-environment growth chambers (Conviron) in the Queen’s University Phytotron at 22 °C with 30% relative humidity and a light intensity of 150 µE m^2^ s^-1^ in short-day (SD) conditions (10 h light/14 h dark cycle). For assessment of stem elongation, the plants designated for phenotyping were transferred to a long-day (LD) growth chamber (14 h light/10 h dark cycle) to enhance the transition from vegetative to reproductive stage, and were photographed at 8 wpg. Plants used for floral dip transformation were grown in LD conditions until bolting. The stems were trimmed one week before the transformation to induce more branches and buds. All soil-grown plants were fertilized every two weeks with 1.5 g L ¹ N:P:K (20:20:20).

When sterile conditions were required, Arabidopsis seeds were incubated in 40% (v/v) bleach for 17 min on a rotating mixer. Seeds were washed 4 times in sterile double-distilled water (ddH_2_O) and plated on MS agar medium (0.5× MSO with vitamins, Cedarlane Labs; 0.8% agar). For selection of transgenic seedlings, kanamycin (50 µg mL ¹) was added to the medium. The seeds were cold-stratified as described above and incubated at room temperature under a light intensity of 50 µE m^2^ s^-1^ for 7–10 days before being transplanted into soil or transferred to liquid MS medium.

*N. benthamiana* plants were grown in an LD chamber (Conviron) under a 16 h light/8 h dark cycle at 22°C with 30% relative humidity and a light intensity of 150 μE m^2^ s^-1^. The plants were fertilized weekly with 1.5 g L ¹ N:P:K (20:20:20).

### Molecular cloning

Clones were generated using various methods as outlined in detail in **Table S1**.

### Phylogenetic analysis

Protein sequences of Arabidopsis CDPKs and CRKs (42 sequences) were retrieved from The Arabidopsis Information Resource (TAIR; Araport11). Predicted proteomes of *Malus × domestica* (GDDH13 v1.1), *Prunus persica* (v2.0.a1), *Rubus occidentalis* (v3.0), and *Fragaria vesca* (v4.0.a2) were obtained from the Genome Database for Rosaceae (GDR) (Jung et al. 2019). Candidate Rosaceae homologs were identified by BLASTP using the Arabidopsis sequences as queries. Candidates were analyzed with HMMER (hmmscan) against the Pfam database, and only proteins containing both a protein kinase domain (PF00069) and at least one EF-hand domain (PF00036, PF13499, or PF13833) were retained. This yielded the Rosaceae CDPKs/CRKs, which together with the 42 Arabidopsis sequences were aligned with MAFFT v7 (Katoh and Standley 2013). A maximum likelihood tree was inferred with IQ-TREE (Nguyen et al. 2015) using the best-fit substitution model selected by ModelFinder (Kalyaanamoorthy et al. 2017). Branch support was assessed using 1,000 ultrafast bootstrap replicates and the SH-aLRT test. Trees were visualized and annotated in iTOL v7.6 (Letunic and Bork 2021). The Subgroup IV clade (**Fig. 1d**) was pruned from the full tree.

**Fig. 1.**
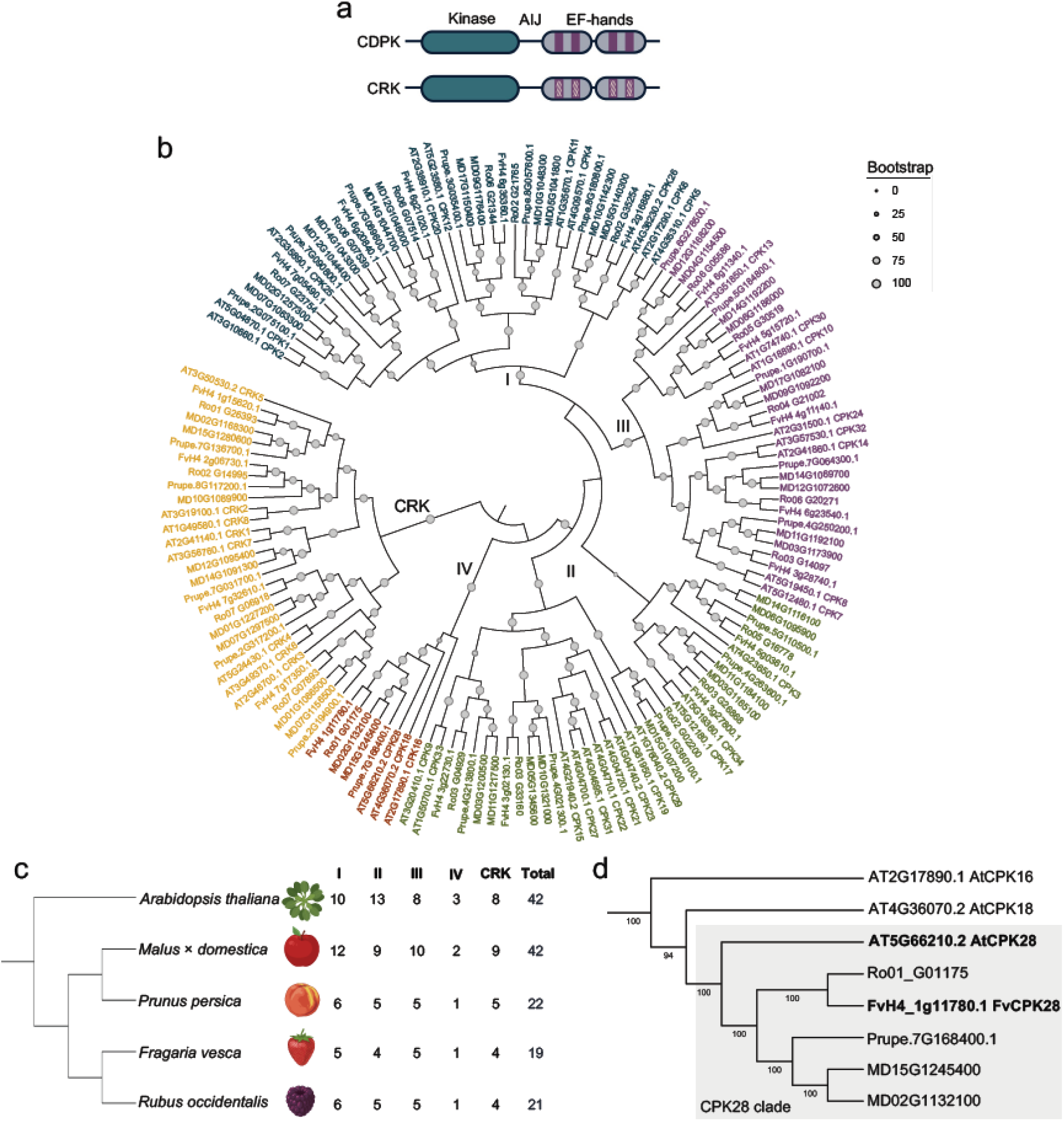
The Rosaceae CDPK/CRK family and identification of CPK28 orthologs. **(a)** Schematic protein architecture of calcium-dependent protein kinases (CDPKs) and CDPK-related kinases (CRKs). Both share a variable N-terminal region, a serine/threonine kinase domain, an autoinhibitory junction (AIJ), and a C-terminal calmodulin-like domain containing four EF-hand motifs. EF-hands are typically functional in CDPKs and degenerate in CRKs (hatched). **(b)** Maximum likelihood phylogeny of CDPKs and CRKs from *Arabidopsis thaliana* (AT), *Malus × domestica* (MD), *Prunus persica* (Prupe), *Fragaria vesca* (Fv), and *Rubus occidentalis* (Ro). The tree is shown rooted on the CRK clade for display. Tip colours denote the five subgroups (I–IV and CRK). Circle size at nodes is proportional to bootstrap support. **(c)** Number of CDPK and CRK genes per species, by subgroup. The cladogram on the left shows the relationships among species and is schematic (branch lengths not to scale; based on NCBI Taxonomy Database). **(d)** Subgroup IV clade pruned from the phylogeny in (b). The shaded box marks the CPK28 clade, comprising *A. thaliana* CPK28 (AtCPK28) and its Rosaceae orthologs. AtCPK28 (At5g66210) and FvCPK28 (FvH4_1g11780.1) are shown in bold. Numbers at nodes indicate bootstrap support (1,000 replicates).

The FvBIK1 sequence reported by (Li et al. 2023) (NCBI accession XM_004307462.2) was mapped to the *F. vesca* v4 genome annotation by BLASTP of its translated protein against the *F. vesca* predicted proteins in GDR, identifying FvH4_7g22540.1 as the corresponding gene. *F. vesca* receptor-like cytoplasmic kinases (RLCKs) were identified by BLASTP using Arabidopsis RLCK subfamily VII sequences as queries, and analyzed as above, retaining sequences containing a protein kinase domain (PF00069 or PF07714). Sequences were aligned with MAFFT and a maximum likelihood tree was inferred with IQ-TREE (1,000 bootstrap replicates), from which the subgroup RLCK VII-8 clade (**Fig. 2b**) was pruned.

**Fig. 2.**
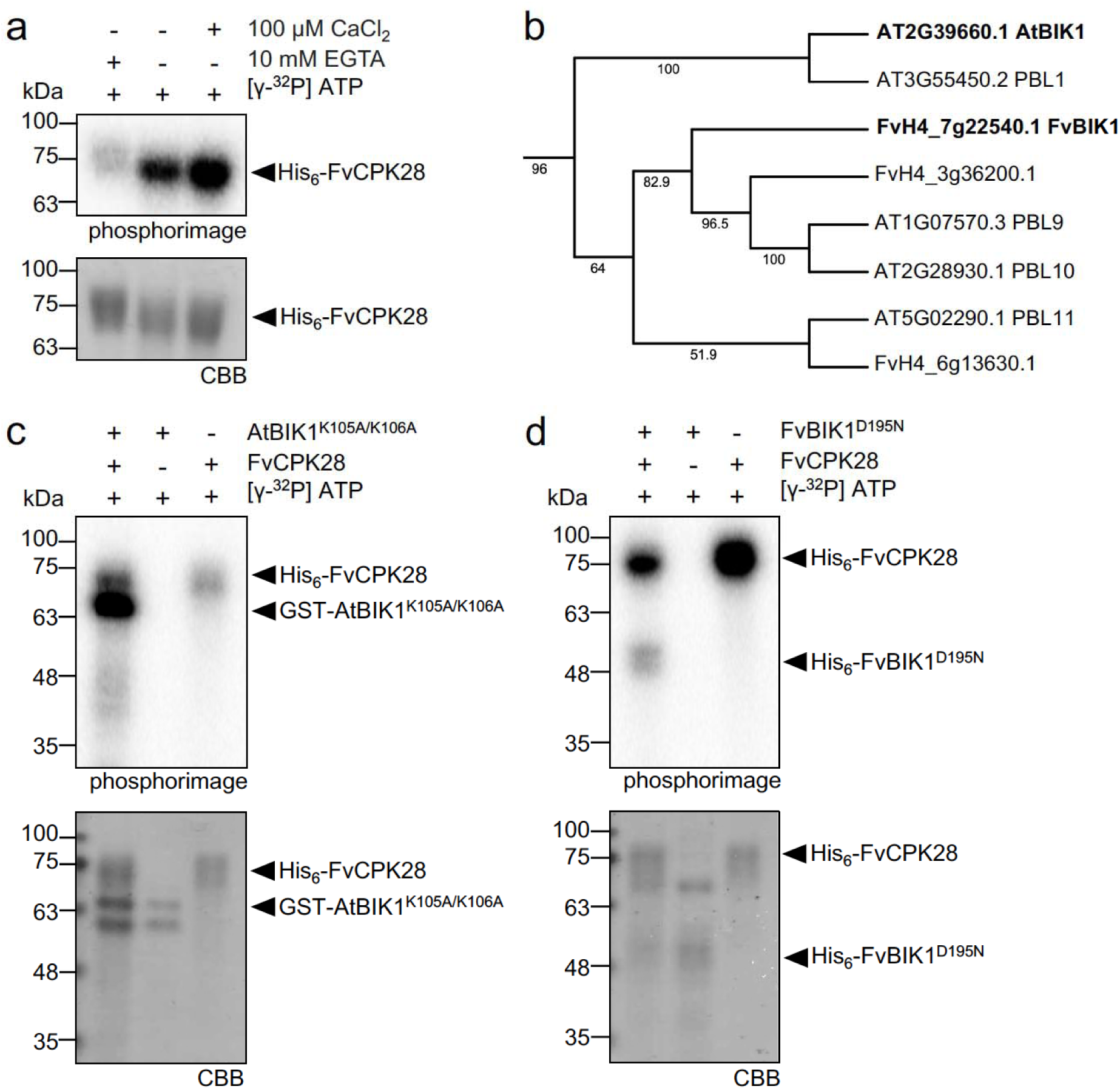
FvCPK28 is a calcium-dependent kinase that phosphorylates AtBIK1 and FvBIK1. **(a)** Autophosphorylation of recombinant His_6_*-*FvCPK28 across a range of free Ca^2+^ concentrations: chelated Ca^2+^ (10 mM EGTA), background Ca^2+^(-EGTA and -CaCl_2_), or with the addition of 100 µM CaCl_2_). **(b)** Maximum likelihood phylogeny of BIK1 and closely related PBL proteins from *Arabidopsis thaliana* and *Fragaria vesca*. FvH4_7g22540.1 is referred to as FvBIK1. AtBIK1 and FvBIK1 are shown in bold. Numbers at nodes indicate bootstrap support. **(c, d)** *In vitro* transphosphorylation of catalytically inactive BIK1 variants by His-FvCPK28. **(c)** GST-AtBIK1^K105A/K106A^. **(d)** His_6_-FvBIK1^D195N^. Each substrate was incubated with or without FvCPK28, and FvCPK28 alone was included as a control. In **(a)**, **(c)** and **(d)**, reactions contained [γ-^32^P]ATP. Proteins were separated by SDS-PAGE, ^32^P incorporation was detected with a phosphorimager and protein loading was assessed by Coomassie Brilliant Blue staining (CBB). Arrowheads indicate the expected positions of the recombinant proteins. All experiments were performed at least three times with similar results

### Recombinant protein expression and purification

All proteins were expressed and purified from *Escherichia coli* strain BL21 using the constructs listed in **Table S1**. Cultures were grown at 37 °C in Luria-Bertani (LB) broth containing appropriate antibiotics until the OD_600_ reached 0.7–0.8. Protein expression was induced with 0.5 mM isopropyl β-D-1-thiogalactopyranoside (IPTG), and cultures were incubated with shaking for 20 h at 25 °C. Cells were harvested by centrifugation at 4,200 × *g* for 15 min at 4 °C. Cell pellets containing His_6_-tagged proteins were resuspended in extraction buffer consisting of 50 mM Tris·HCl (pH 7.5), 100 mM NaCl, and 1 mM phenylmethylsulfonyl fluoride (PMSF). Cell pellets containing the GST-tagged protein were resuspended in phosphate-buffered saline (PBS; Thermo Fisher Scientific) with 1 mM dithiothreitol (DTT) and 1 mM PMSF. Cells were lysed with 1 mg mL ¹ lysozyme for 15 min on ice followed by sonication, as described by (Liese et al. 2024). Lysates were clarified by centrifugation at 15,400 × *g* for 40 min at 4 °C. The supernatants were incubated with either nickel-nitrilotriacetic acid resin (Ni-NTA) (HisPur™ Ni-NTA, Thermo Fisher Scientific) or pre-hydrated glutathione agarose beads (G4510, Sigma-Aldrich) in a conical tube with gentle rotation for 1–2 h at 4 °C. His_6_-tagged proteins were washed sequentially with extraction buffer containing 30 mM and then 40 mM imidazole, and proteins were eluted with extraction buffer containing 500 mM imidazole. The GST-tagged protein was eluted with elution buffer containing 50 mM Tris⋅HCl pH 7.5, 10 mM reduced glutathione and 5 mM DTT. Protein concentrations were determined using Bradford reagent (23200, Thermo Fisher Scientific) with bovine serum albumin (BSA) as the standard. Protein purity was assessed by SDS-PAGE followed by Coomassie Brilliant Blue (CBB) staining. Aliquots were flash-frozen in liquid N_2_ and stored at −80°C until use.

### *In vitro* kinase assays

Autophosphorylation assays were performed by incubating 2 μg of purified His_6_-FvCPK28 in 25 μL of buffer containing 50 mM Tris-HCl (pH 8.0), 25 mM MgCl_2_, 25 mM MnCl_2_, 5 mM DTT, 100 μM ATP, 0.5 μCi [γ-^32^P]ATP and 100 μM CaCl_2_ or 10 mM EGTA.

Trans-phosphorylation assays used 2 μg His_6_-FvCPK28 as the kinase and 4 μg of kinase-dead variants GST-AtBIK1^K105A/K106A^ or His_6_-FvBIK1^D195N^ as the substrates in the same buffer. For each substrate, kinase-alone and substrate-alone reactions were run in parallel as controls. Reactions were kept at room temperature for 30 min, stopped by the addition of 5× Laemmli sample buffer (LSB) and heated at 80 °C for 5 min. Proteins were separated in a 10% SDS-PAGE gel at 80 V for 30 min followed by 150 V for 1 h, in 1× SDS running buffer (25 mM Tris-HCl pH 6.8, 190 mM glycine, 0.1% (w/v) SDS). The gels were sandwiched between two sheets of transparency film, exposed overnight to a storage phosphor screen (Molecular Dynamics). The results were visualized using a Sapphire FL Biomolecular Imager (Azure Biosystems). Gels were post-stained with CBB R-250 (MP Biomedicals) and scanned using an HP OfficeJet Pro 8620. Assays were performed 3 times with similar results.

### *Agrobacterium*-mediated transient expression in *N. benthamiana* and stable transformation of *Arabidopsis*

Binary vectors were transformed into *Agrobacterium tumefaciens* strain GV3101 by electroporation, and transient expression in leaves of 4–5-week-old *N. benthamiana* plants was performed as described previously (Gonçalves Dias et al. 2024). Leaves were analyzed for split-luciferase complementation 3 days post-infiltration (dpi).

Stable transgenic *cpk28-1/35S:FvCPK28-GFP* lines were generated by the floral dip method (Clough and Bent 1998). Transformants were selected through multiple generations to homozygosity on MS agar medium containing kanamycin (50 µg mL ¹) and used for further analysis.

### Split-luciferase complementation assay (SLCA)

Split-luciferase complementation assays were performed using *A. tumefaciens* GV3101 strains carrying *pCAMBIA1300-nLuc/cLuc* or *pGWB-nLuc/cLuc* constructs (**Table S1**), with two complementary approaches in parallel. For quantification, the assay was performed as described previously (Gonçalves Dias et al. 2024), and luminescence was measured with a SpectraMax Paradigm microplate reader (Molecular Devices). For luminescence imaging, the assay was performed according to (Wang et al. 2020, 2021). Luminescence was captured with a cooled CCD imaging system (NightSHADE LB 985, Berthold Technologies) and analyzed with indiGO software. Both assays were repeated at least five times independently with similar results.

### Confocal microscopy

Images from 5–8-day-old *cpk28-1/35S:FvCPK28-GFP* roots were taken using a Zeiss LSM 980 confocal microscope equipped with an Airyscan 2 detector on an inverted Axio Observer 7 stand (Carl Zeiss, Jena, Germany) in the Biology Department at Queen’s University. Roots were stained with 2 μM FM464 (Calbiochem) before mounting in water on a glass slide. GFP and FM4-64 were excited sequentially with 488 nm and 561 nm lasers, respectively, and emission was collected at 499–548 nm (GFP) and 573–627 nm (FM4-64). Images were acquired in Fast Airyscan mode and processed using Airyscan processing in ZEN 3.13 (Carl Zeiss).

### Immune assays

Immunogenic elf18 and AtPep1 peptides used for immune assays were synthesized by EZ Biotech (Indiana, USA). Immune-induced ROS production and seedling growth inhibition assays were performed on 4- to 5-week-old soil-grown plants or 12-day-old seedlings, respectively, as previously described (Bredow et al. 2019).

### Statistics

GraphPad Prism 11 was used to perform statistical tests on all quantitative data.

## Results

### The CDPK/CRK family in Rosaceae

CDPKs and CRKs consist of a variable N-terminal domain, a serine/threonine protein kinase domain, and a C-terminal calmodulin-like domain, which are linked by an auto-inhibitory junction. The calmodulin-like domain contains EF-hand motifs that are typically functional in CDPKs but degenerate in CRKs (**Fig. 1a**) (Bredow and Monaghan 2019; Yip Delormel and Boudsocq 2019).

To place the Rosaceae CDPK/CRK family in a phylogenetic context, we constructed a maximum likelihood phylogeny of CDPKs and CRKs from Arabidopsis and four Rosaceae species (*Malus* × *domestica*, *Prunus persica*, *Rubus occidentalis*, and *Fragaria vesca*) (**Fig. 1b**). Homologs were identified in each genome using the 42 *Arabidopsis* sequences as queries and retaining proteins with both a kinase and EF-hand domain, yielding 104 Rosaceae CDPKs/CRKs (146 sequences in total; **Fig. 1c**). The proteins clustered into the five recognized subgroups: CDPK subgroups I–IV and the CRK clade (Yip Delormel and Boudsocq 2019), each containing members from all five species. Subgroups I, II, and III were the largest, while subgroup IV, which includes *Arabidopsis* CPK28, was the smallest, with only 8 members across all species (**Fig. 1c**).

To identify CPK28 orthologs in Rosaceae, we examined the Subgroup IV clade of the phylogeny (**Fig. 1d**). Each Rosaceae genome encoded one (*F. vesca*, *R. occidentalis*, *P. persica*) or two (*M.* × *domestica*) Subgroup IV members, all of which grouped with AtCPK28 in a well-supported clade (bootstrap 100) distinct from the Brassicaceae-specific Subgroup IV CDPKs AtCPK16 and AtCPK18 (Dou et al. 2026) (**Fig. 1d**), identifying them as CPK28 orthologs. Whether these orthologs share the biochemical and immune functions of AtCPK28 has not been tested. We therefore selected the single *F. vesca* ortholog (FvH4_1g11780.1, hereafter FvCPK28) as a representative to address this question.

### FvCPK28 is a Ca^2+^-dependent kinase that phosphorylates AtBIK1 and FvBIK1

CPK28 orthologs characterized to date are Ca^2+^-activated kinases (Bender et al. 2017; Wang et al. 2018a; Bredow et al. 2021; Hu et al. 2021; Dou et al. 2026). To test whether FvCPK28 shares this biochemical property, we assessed the autophosphorylation of purified recombinant FvCPK28 N-terminally tagged with six tandem histidine residues (His_6_-FvCPK28) under varying Ca^2+^ conditions using [γ-^32^P]ATP. In the presence of the Ca^2+^ chelator EGTA (10 mM), His_6_-FvCPK28 showed weak but detectable autophosphorylation, which increased substantially under background Ca^2+^ conditions (no EGTA or added CaCl_2_) and was further enhanced by the addition of 100 µM CaCl_2_ (**Fig. 2a**). Thus, recombinant FvCPK28 is catalytically active *in vitro*, and its autophosphorylation is responsive to Ca^2+^, indicating that it is a functional Ca^2+^-dependent protein kinase.

In Arabidopsis, CPK28 regulates the accumulation of BIK1 to modulate the amplitude of PTI (Monaghan et al. 2014). In strawberry, recent work showed that FvBIK1 is induced during brassinosteroid-elicited resistance of fruit to *Botrytis cinerea* and interacts with FvBAK1 and FvRBOHD in yeast two-hybrid assays (Li et al. 2023).

Using the NCBI accession reported for FvBIK1 (Li et al. 2023), we identified the corresponding gene in the *F. vesca* genome (FvH4_7g22540.1) and performed a phylogenetic analysis of the RLCK subgroup VII-8 members AtBIK1, AtPBL1, AtPBL9, AtPBL10, and AtPBL11, together with their *F. vesca* homologs (**Fig. 2b**). AtBIK1 grouped with AtPBL1, whereas FvBIK1 grouped with AtPBL9 and AtPBL10. Like AtBIK1, AtPBL9 and AtPBL10 are transcriptionally induced by multiple molecular patterns (Rao et al. 2018), consistent with a potential role for this clade in immune signaling. Importantly, FvBIK1 interacts in yeast with FvBAK1 and FvRBOHD (Li et al. 2023), putative strawberry homologs of core BIK1 partners in Arabidopsis. We therefore used FvBIK1 as a candidate substrate for FvCPK28.

We first asked whether FvCPK28 can phosphorylate AtBIK1, a previously characterized CPK28 substrate (Monaghan et al. 2014). To distinguish trans-phosphorylation from substrate autophosphorylation, we used the kinase-inactive variant GST-AtBIK1^K105A/K106A^ (Bredow et al. 2021). When incubated alone, AtBIK1^K105A/K106A^ showed no detectable ^32^P incorporation, whereas co-incubation with His-FvCPK28 resulted in strong phosphorylation of AtBIK1^K105A/K106A^ (**Fig. 2c**), indicating that FvCPK28 directly phosphorylates AtBIK1 *in vitro*.

We next tested if FvCPK28 can phosphorylate FvBIK1. Because no kinase-inactive variant of FvBIK1 was available, we identified Asp195 as the catalytic aspartate of the HRD motif (equivalent to AtBIK1 Asp202 (Mühlenbeck et al. 2024) and substituted it with asparagine (FvBIK1^D195N^), an isosteric substitution that preserves native side-chain interactions while abolishing catalytic activity (Reinhardt and Leonard 2023). When incubated alone, His_6_-FvBIK1^D195N^ showed no detectable ^32^P incorporation, confirming the loss of autophosphorylation, whereas co-incubation with His_6_-FvCPK28 resulted in phosphorylation of FvBIK1^D195N^ (**Fig. 2d**). Together, these results show that FvCPK28 can directly phosphorylate both AtBIK1 and FvBIK1 *in vitro*.

### FvCPK28 associates with AtBIK1 and FvBIK1 *in planta*

Having shown that FvCPK28 phosphorylates AtBIK1 and FvBIK1 *in vitro*, we next asked whether FvCPK28 associates with these proteins *in vivo*. We performed split-luciferase complementation assays in *N. benthamiana*, co-expressing FvCPK28-nLuc with cLuc-AtBIK1 or cLuc-FvBIK1, compared to co-expression with WALL-ASSOCIATED KINASE 1 (WAK1)-nLuc as a negative control. Co-expression of FvCPK28-nLuc with either c-LucAtBIK1 or cLuc-FvBIK1 reconstituted luciferase activity significantly above the corresponding WAK1-nLuc controls (**Fig. 3a, b**), with the strongest signal arising from the FvCPK28–AtBIK1 pair (**Fig. 3b**). Thus, in addition to phosphorylating both proteins *in vitro*, FvCPK28 can associate with AtBIK1 and FvBIK1 *in planta*.

**Fig. 3.**
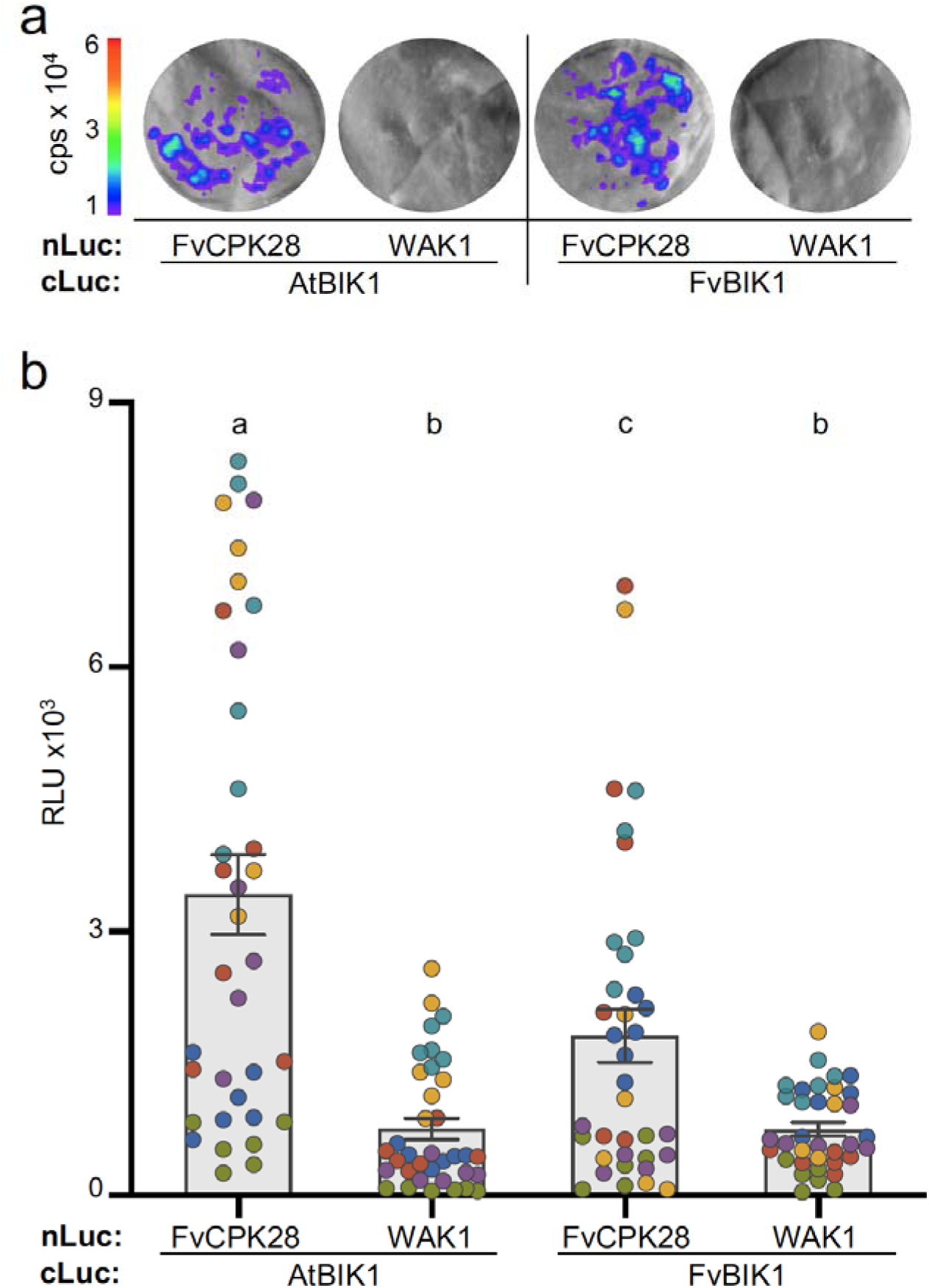
FvCPK28 associates with AtBIK1 and FvBIK1 *in vivo*. Split-luciferase complementation assays with FvCPK28-nLuc or AtWAK1-nLuc with cLuc-AtBIK1 or cLuc-FvBIK1, transiently co-expressed in *Nicotiana benthamiana*. **(a)** Representative luminescence images of the infiltrated leaf area, captured with a cooled CCD imaging system (NightSHADE LB 985). The colour scale indicates luminescence intensity (counts per second, cps ×10). **(b)** Luminescence quantified from leaf discs using a microplate reader (SpectraMax Paradigm), expressed as relative luminescence units (RLU ×10³). Each dot represents one leaf disc from six independent experiments (n=6), shown in different colours. Bars indicate the mean ± SEM, and different letters indicate statistically significant differences (one-way ANOVA followed by Tukey’s post hoc test, *P* < 0.0001)

### FvCPK28 complements the developmental and immune phenotypes of the Arabidopsis *cpk28-1* mutant

Having established that FvCPK28 is an active Ca^2+^-dependent kinase that phosphorylates and associates with AtBIK1 and FvBIK1, we asked whether it can functionally replace AtCPK28 *in planta*. We generated transgenic lines expressing FvCPK28 C-terminally tagged with green fluorescent protein (GFP) under the CaMV *35S* promoter in the Arabidopsis loss-of-function mutant *cpk28-1* (*cpk28-1/35S:FvCPK28-GFP*). We analyzed two independent homozygous lines (#13 and #14) alongside wild-type Col-0, *cpk28-1*, and *cpk28-1* complemented with AtCPK28 (*cpk28-1/35S:AtCPK28-YFP*) as a positive control. Like AtCPK28-YFP (Monaghan et al. 2014), FvCPK28-GFP localized to the plasma membrane of root epidermal cells, as shown by co-localization with the membrane dye FM4-64 (**Fig. 4a**).

**Fig. 4.**
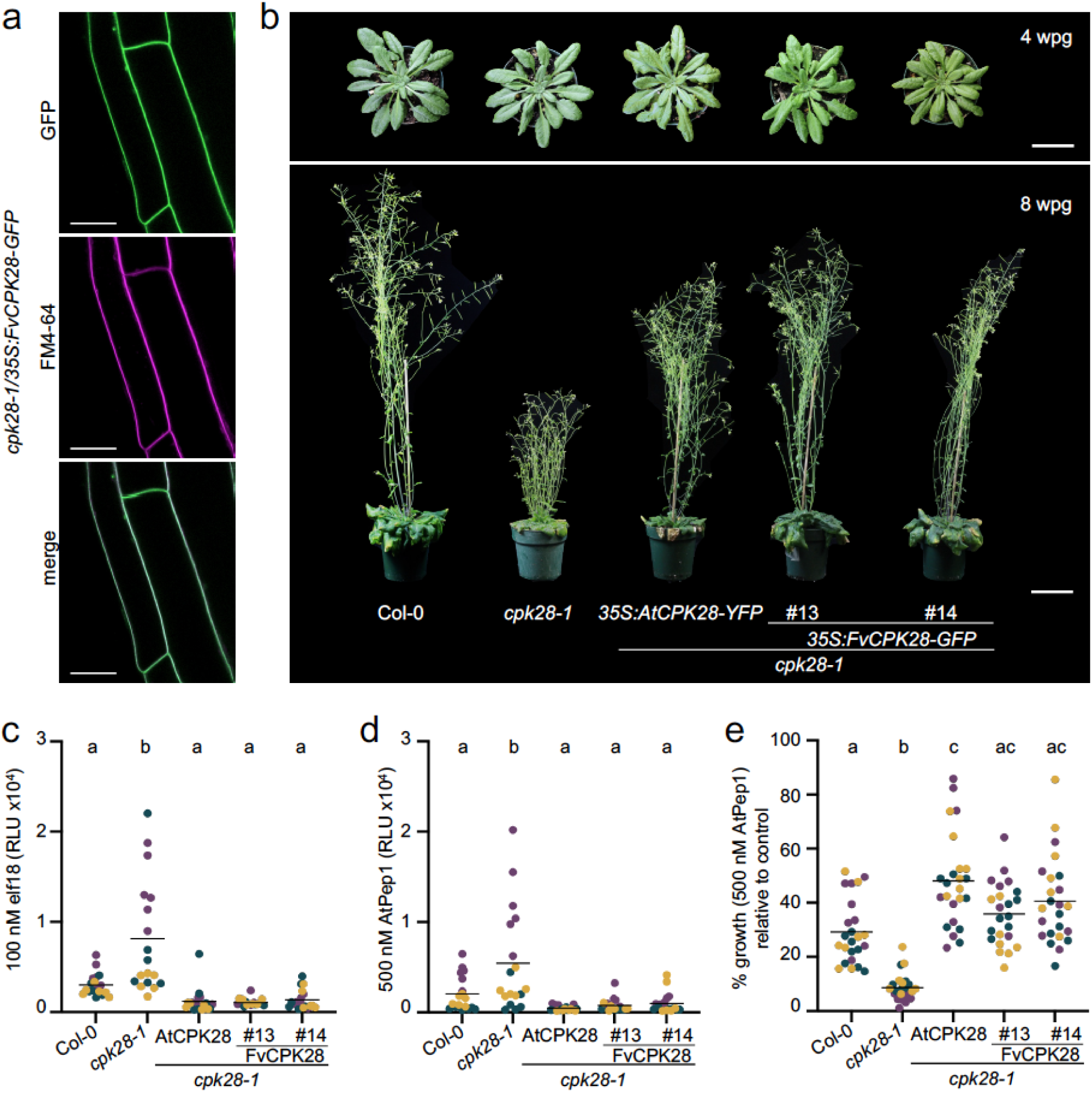
FvCPK28 functionally complements the *Arabidopsis cpk28-1* mutant. **(a)** Confocal micrographs of root cells from *cpk28-1/35S:FvCPK28-GFP* line 14 indicating the subcellular localization of FvCPK28-GFP (GFP) and the plasma membrane (FM4-64) and both images merged. Scale bar: 20 μm. **(b)** Morphology of Col-0 (wild type), *cpk28-1*, and *cpk28-1* plants expressing *35S:AtCPK28-YFP* or *35S:FvCPK28-GFP* (independent lines #13 and #14). Upper: rosettes at 4 weeks post-germination (wpg) under short-day conditions. Lower: the same genotypes at 8 wpg after transfer to long-day conditions, showing stem elongation. Images are representative of 6 plants per genotype. Scale bars = 2 cm (upper) and 5 cm (lower). **(c, d)** Total reactive oxygen species (ROS) production in leaf discs treated with 100 nM elf18 **(c)** or 500 nM AtPep1 **(d)**, expressed as relative light units (RLU ×10). **(e)** Seedling growth inhibition (SGI) in response to 500 nM AtPep1, expressed as percentage growth relative to untreated controls. In (**c–e**), each dot represents one biological replicate (n = 6 per genotype per experiment in c and d; n = 8 in e), colours denote three independent experiments, and horizontal bars indicate the mean. Different letters indicate statistically significant differences (one-way ANOVA followed by Tukey’s post hoc test, (P < 0.001)

Following the transition to the reproductive stage, *cpk28-1* mutants exhibit developmental phenotypes, including shorter leaf petioles, enhanced anthocyanin accumulation, and reduced stem elongation (Matschi et al. 2013, 2015). Both *cpk28-1/35S:FvCPK28-GFP* lines restored stem elongation to wild-type levels, comparable to the AtCPK28-complemented control (**Fig. 4b**), indicating that FvCPK28 rescues the developmental defect caused by loss of AtCPK28.

In addition to these developmental defects, loss of CPK28 function results in enhanced immune responses (Monaghan et al. 2014). CPK28 negatively regulates early immune signaling by controlling BIK1 turnover (Wang et al. 2018a; Dou et al. 2025). Consequently, *cpk28* loss-of-function mutants accumulate elevated levels of BIK1 and produce a heightened apoplastic reactive oxygen species (ROS) burst in response to immunogenic peptides (Monaghan et al. 2014). Consistent with this, *cpk28-1* produced a heightened ROS burst relative to Col-0 in response to both the bacterial elicitor elf18 (the N-terminal 18 residues of elongation factor Tu) and the endogenous peptide AtPep1 (**Fig. 4c–d**). Both *cpk28-1/35S:FvCPK28-GFP* lines restored the response to wild-type levels, as did the *cpk28-1/35S:AtCPK28-YFP* control (**Fig. 4c–d**), demonstrating that FvCPK28 can functionally replace AtCPK28 in early immune responses.

To assess whether FvCPK28 also restores a sustained immune output, we measured elicitor-induced seedling growth inhibition (SGI), which is enhanced in *cpk28-1* owing to its heightened immune signaling (Monaghan et al. 2014). Following treatment with AtPep1, *cpk28-1* seedlings showed greater growth inhibition than Col-0, while both *cpk28-1/35S:FvCPK28-GFP* lines showed reduced sensitivity comparable to wild type and to the AtCPK28 control (**Fig. 4e**). Together, these results demonstrate that FvCPK28 functionally complements the loss of AtCPK28 across a developmental phenotype and both early (ROS) and sustained (SGI) immune signaling outputs, establishing that the strawberry ortholog retains CPK28 function in Arabidopsis. Similar functional conservation of CPK28 orthologs has been reported in other plant species, such as rice, cotton, and tomato (Wang et al. 2018b; Wu et al. 2021; Ding et al. 2022), suggesting that the role of this kinase in modulating immune signaling is likely conserved across angiosperms.

## Discussion

Calcium-dependent protein kinases translate cellular Ca^2+^ signals into phosphorylation events that shape a broad range of cellular processes, including plant growth and immunity (Yip Delormel and Boudsocq 2019). Among them, CPK28 acts as a negative regulator of immune signaling while also contributing to the vegetative-to-reproductive stage transition and other pathways (Matschi et al. 2013; Monaghan et al. 2014; Jin et al. 2017; Hu et al. 2021). In Rosaceae, the CDPK family has been catalogued across several species (Wei et al. 2016; Xiong et al. 2022), and expression of apple *MdCPK28* was found to be significantly down-regulated in response to infection by *Alternaria alternata* (Wei et al. 2016), suggesting a possible role in modulating host immunity. However, whether a Rosaceae CPK28 ortholog functions in plant immunity has not been tested. Here, we used updated genome assemblies to identify CPK28 orthologs in Rosaceae and selected the *Fragaria vesca* ortholog, FvCPK28, as a case study to ask whether CPK28 function is conserved in this lineage.

Our phylogenetic reconstruction of the CDPK/CRK family across four Rosaceae species and Arabidopsis recovered the conserved family structure of four CDPK subgroups and the CRK clade (Yip Delormel and Boudsocq 2019). Unlike earlier surveys of Rosaceae, which focused on CDPKs (Wei et al. 2016; Xiong et al. 2022), our analysis additionally resolved the CRK clade. Consistent with these earlier surveys, Subgroup IV, which contains CPK28, was the smallest clade, and all Rosaceae Subgroup IV members formed a well-supported clade with AtCPK28. These results indicate that CPK28 orthologs are present in all four Rosaceae species examined, forming a distinct lineage with AtCPK28.

FvCPK28 retains the defining biochemical properties of AtCPK28. Its autophosphorylation was reduced in the presence of EGTA and strongly stimulated by Ca²□, as described for AtCPK28 and the rice ortholog OsCPK4 (Matschi et al. 2013; Bender et al. 2017; Bredow et al. 2021). In Arabidopsis, CPK28 buffers immune signaling by limiting the accumulation of BIK1, a rate-limiting component of PTI (Monaghan et al. 2014; Wang et al. 2018a). FvCPK28 directly phosphorylated both AtBIK1 and FvBIK1 *in vitro* and associated with both proteins *in planta*, suggesting that the CPK28–BIK1 kinase–substrate relationship is retained in the strawberry orthologs. FvBIK1 does not group directly with AtBIK1, and RLCK subfamily VII comprises many closely related kinases with diverse functions (Rao et al. 2018). Whether FvBIK1 is the main CPK28 substrate in strawberry therefore remains to be determined.

The ability of FvCPK28 to phosphorylate and associate with AtBIK1 provides a plausible mechanistic basis for its function in Arabidopsis. Because obtaining stable transgenic or genome-edited strawberry lines takes several months (Zhou et al. 2018; Wilson et al. 2019), which restricts such approaches to a few candidate genes (Guidarelli and Baraldi 2015), we used the well-characterized Arabidopsis *cpk28-1* mutant to test FvCPK28 function *in planta*. Like AtCPK28, FvCPK28 localized to the plasma membrane and complemented both the developmental and immune phenotypes of *cpk28-1*, restoring stem elongation as well as early (elicitor-induced ROS) and sustained (SGI) immune outputs. Thus, FvCPK28 can functionally replace AtCPK28 in Arabidopsis.

We acknowledge that these experiments assess the behaviour of FvCPK28 within the Arabidopsis system, and they establish that the strawberry protein can complement AtCPK28 function in this context, but do not by themselves define how FvCPK28 acts in strawberry. FvCPK28 was expressed under the constitutive *35S* promoter in a heterologous system, and its association with BIK1 orthologs was tested by transient expression in *N. benthamiana*. Establishing whether FvCPK28 regulates immune homeostasis in strawberry will require loss-of-function or overexpression lines in *F. vesca*, generated by tools such as CRISPR/Cas9 or RNAi (Guidarelli and Baraldi 2015; Zhou et al. 2018). This approach has proven informative in other crops, where CPK28 orthologs were characterized directly in their native species and shown to negatively regulate immunity, including rice OsCPK4, cotton GhCDPK28, and tomato SlCPK28 (Wang et al. 2018b; Wu et al. 2021; Ding et al. 2022). Together with these studies, our results suggest that the immune-regulatory function of CPK28 is conserved in Rosaceae.

As a multifunctional kinase, CPK28 contributes to diverse physiological processes, ranging from vegetative-to-reproductive phase transition (Matschi et al. 2013) to abiotic stress signaling (Jin et al. 2017; Hu et al. 2021) and immunity (Monaghan et al. 2014). This position at the intersection of development and immune homeostasis makes CPK28 an attractive target for breeding strategies aimed at optimizing growth–defense trade-offs. For example, in pear (*Pyrus bretschneideri*), PbrCPK28 has been associated with fructose accumulation in developing fruit (Wu et al. 2023), suggesting that CPK28 orthologs may influence fruit quality in Rosaceae. Together with the conservation of its immune-buffering function shown here, CPK28 emerges as a promising candidate for modulating traits at the intersection of immunity, development, and fruit quality in Rosaceae crops.

Taken together, our results identify CPK28 orthologs across Rosaceae and show, using FvCPK28 as a case study, that the strawberry ortholog is a Ca²-dependent kinase that phosphorylates and associates with BIK1 orthologs and complements the developmental and immune phenotypes of Arabidopsis *cpk28-1*. Given the role of CPK28 in immune optimization, FvCPK28 provides an entry point for future efforts to balance immunity and growth in strawberry and related crops.

## Acknowledgements

We are grateful to all members of our lab, past and present, for engaging discussions over the course of this project, and for reviewing our manuscript before submission. We thank Dr. Beatriz Appezzato da Glória (University of São Paulo) for her support and guidance in the early stages of this research. Queen’s University is situated on the territory of the Haudenosaunee and Anishinaabek, and we are grateful to live, work, and play on these lands.

## Author Contributions

M.G.D.: Conceptualization; data curation; formal analysis; funding acquisition; investigation; visualization; writing – original draft; writing – review & editing.

J.M.: Conceptualization; funding acquisition; supervision; writing – review & editing.

## Data Availability

Any materials described in this article will be made freely available upon request.

## References

Bender KW, Blackburn RK, Monaghan J, et al (2017) Autophosphorylation-based Calcium (Ca2+) Sensitivity Priming and Ca2+/Calmodulin Inhibition of Arabidopsis thaliana Ca2+-dependent Protein Kinase 28 (CPK28). J Biol Chem 292:3988–4002

Bredow M, Bender KW, Johnson Dingee A, et al (2021) Phosphorylation-dependent subfunctionalization of the calcium-dependent protein kinase CPK28. Proc Natl Acad Sci U S A 118.: 10.1073/pnas.2024272118

Bredow M, Monaghan J (2019) Regulation of Plant Immune Signaling by Calcium-Dependent Protein Kinases. Mol Plant Microbe Interact 32:6–19

Bredow M, Sementchoukova I, Siegel K, Monaghan J (2019) Pattern-Triggered Oxidative Burst and Seedling Growth Inhibition Assays in Arabidopsis thaliana. J Vis Exp. 10.3791/59437

Chinchilla D, Zipfel C, Robatzek S, et al (2007) A flagellin-induced complex of the receptor FLS2 and BAK1 initiates plant defence. Nature 448:497–500

Clough SJ, Bent AF (1998) Floral dip: a simplified method for Agrobacterium-mediated transformation of Arabidopsis thaliana. Plant J 16:735–743

Crizel RL, Perin EC, Vighi IL, et al (2020) Genome-wide identification, and characterization of the CDPK gene family reveal their involvement in abiotic stress response in Fragaria x ananassa. Sci Rep 10:11040

Da Costa Arruda GC, Kovács M, Pomázi A (2024) Biological control mechanisms of non Saccharomyces yeasts against filamentous fungal pathogens within Rosaceae family. Int J Agron 2024.: 10.1155/2024/1538220

DeFalco TA, Zipfel C (2021) Molecular mechanisms of early plant pattern-triggered immune signaling. Mol Cell 81:4346

Ding Y, Yang H, Wu S, et al (2022) CPK28-NLP7 module integrates cold-induced Ca2+ signal and transcriptional reprogramming in Arabidopsis. Sci Adv 8:eabn7901

Dong H, Wu C, Luo C, Wei M, Qu S, Wang S (2020) Overexpression of MdCPK1a gene, a calcium dependent protein kinase in apple, increase tobacco cold tolerance via scavenging ROS accumulation. PLoS ONE 15:e0242139

Dou R, Castel B, Tanney CAS, et al (2026) Divergent roles of CPK28 in immune homeostasis across land plants. Mol Plant Microbe Interact 39:573–579

Dou R, Miguel VN, Grubb LE, et al (2025) Class IV plant U-box proteins function redundantly to optimize protein accumulation of receptor-like cytoplasmic kinase BIK1. Plant Physiol 198:kiaf267

Gonçalves Dias M, Doss B, Rawat A, et al (2024) Subfamily C7 Raf-like kinases MRK1, RAF26, and RAF39 regulate immune homeostasis and stomatal opening in Arabidopsis thaliana. New Phytol 244:2278–2294

Gonçalves Dias M, Soleimani F, Monaghan J (2022) Activation and turnover of the plant immune signaling kinase BIK1: a fine balance. Essays Biochem 66:207–218

Guidarelli M, Baraldi E (2015) Transient transformation meets gene function discovery: the strawberry fruit case. Front Plant Sci 6:444

Heese A, Hann DR, Gimenez-Ibanez S, et al (2007) The receptor-like kinase SERK3/BAK1 is a central regulator of innate immunity in plants. Proc Natl Acad Sci U S A 104:12217–12222

Hrabak EM, Chan CWM, Gribskov M, et al (2003) The Arabidopsis CDPK-SnRK superfamily of protein kinases. Plant Physiol 132:666–680

Huot B, Yao J, Montgomery BL, He SY (2014) Growth-defense tradeoffs in plants: a balancing act to optimize fitness. Mol Plant 7:1267–1287

Hu Z, Li J, Ding S, et al (2021) The protein kinase CPK28 phosphorylates ascorbate peroxidase and enhances thermotolerance in tomato. Plant Physiology 186:1302–1317

Jin Y, Ye N, Zhu F, et al (2017) Calcium-dependent protein kinase CPK28 targets the methionine adenosyltransferases for degradation by the 26S proteasome and affects ethylene biosynthesis and lignin deposition in Arabidopsis. Plant J 90:304–318

Jung S, Lee T, Cheng C-H, et al (2019) 15 years of GDR: New data and functionality in the Genome Database for Rosaceae. Nucleic Acids Res 47:D1137–D1145

Kadota Y, Sklenar J, Derbyshire P, et al (2014) Direct regulation of the NADPH oxidase RBOHD by the PRR-associated kinase BIK1 during plant immunity. Mol Cell 54:43–55

Kalyaanamoorthy S, Minh BQ, Wong TKF, et al (2017) ModelFinder: fast model selection for accurate phylogenetic estimates. Nat Methods 14:587–589

Katoh K, Standley DM (2013) MAFFT multiple sequence alignment software version 7: improvements in performance and usability. Mol Biol Evol 30:772–780

Lallemand T, Leduc M, Desmazières A, et al (2023) Insights into the evolution of ohnologous sequences and their epigenetic marks post-WGD in Malus domestica. Genome Biol Evol 15:evad178

Letunic I, Bork P (2021) Interactive Tree Of Life (iTOL) v5: an online tool for phylogenetic tree display and annotation. Nucleic Acids Res 49:W293–W296

Li C, Wang K, Tan M, et al (2023) Involvement of a receptor-like kinase complex of FvFLS2 and FvBAK1 in brassinosteroids-induced immunity in postharvest strawberry fruit. Postharvest Biol Technol 198:112266

Liese A, Eichstädt B, Lederer S, et al (2024) Imaging of plant calcium-sensor kinase conformation monitors real time calcium-dependent decoding in planta. Plant Cell 36:276– 297

Li L, Li M, Yu L, et al (2014) The FLS2-associated kinase BIK1 directly phosphorylates the NADPH oxidase RbohD to control plant immunity. Cell Host Microbe 15:329–338

Lu D, Wu S, Gao X, et al (2010) A receptor-like cytoplasmic kinase, BIK1, associates with a flagellin receptor complex to initiate plant innate immunity. Proc Natl Acad Sci U S A 107:496–501

Matschi S, Hake K, Herde M, et al (2015) The calcium-dependent protein kinase CPK28 regulates development by inducing growth phase-specific, spatially restricted alterations in jasmonic acid levels independent of defense responses in Arabidopsis. Plant Cell 27:591– 606

Matschi S, Werner S, Schulze WX, et al (2013) Function of calcium-dependent protein kinase CPK28 of Arabidopsis thaliana in plant stem elongation and vascular development. Plant J 73:883–896

Monaghan J, Matschi S, Shorinola O, et al (2014) The calcium-dependent protein kinase CPK28 buffers plant immunity and regulates BIK1 turnover. Cell Host Microbe 16:605–615

Mühlenbeck H, Tsutsui Y, Lemmon MA, et al (2024) Allosteric activation of the co-receptor BAK1 by the EFR receptor kinase initiates immune signaling. Elife 12:RP92110

Nguyen L-T, Schmidt HA, von Haeseler A, Minh BQ (2015) IQ-TREE: a fast and effective stochastic algorithm for estimating maximum-likelihood phylogenies. Mol Biol Evol 32:268– 274

Rao S, Zhou Z, Miao P, et al (2018) Roles of Receptor-Like Cytoplasmic Kinase VII Members in Pattern-Triggered Immune Signaling. Plant Physiol 177:1679–1690

Reinhardt R, Leonard TA (2023) A critical evaluation of protein kinase regulation by activation loop autophosphorylation. Elife 12:e88210

Shi J, Ma Y, Wang D, Wang F (2025) MdCDPK24 encoding calcium-dependent protein kinase enhances apple resistance to Colletotrichum gloeosporioides. Horticulturae 11:942

Shulaev V, Sargent DJ, Crowhurst RN, et al (2011) The genome of woodland strawberry (Fragaria vesca). Nat Genet 43:109–116

Thor K, Jiang S, Michard E, et al (2020) The calcium-permeable channel OSCA1.3 regulates plant stomatal immunity. Nature 585:569–573

Valmonte GR, Arthur K, Higgins CM, MacDiarmid RM (2014) Calcium-dependent protein kinases in plants: evolution, expression and function. Plant Cell Physiol 55:551–569

Wang J, Grubb LE, Wang J, et al (2018a) A regulatory module controlling homeostasis of a plant immune kinase. Mol Cell 69:493–504.e6

Wang J, Wang S, Hu K, et al (2018b) The Kinase OsCPK4 Regulates a Buffering Mechanism That Fine-Tunes Innate Immunity. Plant Physiol 176:1835–1849

Wang L, Ding Y, He L, et al (2020) A virus-encoded protein suppresses methylation of the viral genome through its interaction with AGO4 in the Cajal body. Elife 9:e55542

Wang L, Yu G, Macho A, Lozano-Durán R (2021) Split-luciferase complementation imaging assay to study protein-protein interactions in Nicotiana benthamiana. Bio Protoc 11.: 10.21769/bioprotoc.4237

Wei M, Wang S, Dong H, et al (2016) Characterization and comparison of the CPK gene family in the apple (Malus × domestica) and other Rosaceae species and its response to Alternaria alternata infection. PLoS One 11:e0155590

Wilson FM, Harrison K, Armitage AD, et al (2019) CRISPR/Cas9-mediated mutagenesis of phytoene desaturase in diploid and octoploid strawberry. Plant Methods 15:45

Wu J, Zhu RX, Li JM, et al (2023) Pear PbrCPK28 gene and application thereof. Chinese Patent No. CN116042650A

Wu Y, Zhang L, Zhou J, et al (2021) Calcium-Dependent Protein Kinase GhCDPK28 Was Dentified and Involved in Verticillium Wilt Resistance in Cotton. Front Plant Sci 12:772649

Xie K, Chen J, Wang Q, Yang Y (2014) Direct phosphorylation and activation of a mitogen-activated protein kinase by a calcium-dependent protein kinase in rice. Plant Cell 26:3077–3089

Xiong Y, Lin D, Ma S, et al (2022) Genome-wide identification of the calcium-dependent protein kinase gene family in Fragaria vesca and expression analysis under different biotic stresses. Eur J Plant Pathol 164:283–298

Yip Delormel T, Boudsocq M (2019) Properties and functions of calcium-dependent protein kinases and their relatives in Arabidopsis thaliana. New Phytol 224:585–604

Zhang J, Li W, Xiang T, et al (2010) Receptor-like Cytoplasmic Kinases Integrate Signaling from Multiple Plant Immune Receptors and Are Targeted by a Pseudomonas syringae Effector. Cell Host Microbe 7:290–301

Zhou J, Wang G, Liu Z (2018) Efficient genome editing of wild strawberry genes, vector development and validation. Plant Biotechnol J 16:1868–1877

